# ARBoR: An Identity and Security Solution for Clinical Reporting

**DOI:** 10.1101/567875

**Authors:** Eric Venner, Mullai Murugan, Walker Hale, Jordan Jones, Shan Lu, Victoria Yi, Richard A. Gibbs

## Abstract

**Motivation:** Clinical genome sequencing laboratories return reports containing clinical testing results, signed by an Board Certified clinical geneticist, to the ordering physician. This report is often a pdf, but can also be a physical paper copy or a structured data file. The reports are frequently modified and re-issued, due to changes in variant interpretation or clinical attributes. To precisely track report authenticity we developed ARBoR, an application for tracking the lineage of versioned clinical reports even when they are distributed as pdf or paper copies. ARBoR employs a modified blockchain approach and instead of relying on a computationally intensive consensus mechanism for determining authenticity, we allow supervised and digitally signed writes to an encrypted ledger, which is then exactly replicated to many clients.

**Results:** ARBoR was implemented for clinical reporting in the HGSC-CL Clinical Laboratory, initially as part of the NIH’s Electronic Medical Record and Genomics (eMERGE) project. To date we have issued 15,205 versioned clinical reports tracked by ARBoR. This system has provided us with a simple and tamper-proof mechanism for tracking clinical reports with a complicated update history.

## BACKGROUND

Clinical genome sequencing has made a major impact on the diagnosis and treatment of a broad range of clinical presentations, notably within the care of cases of pediatric cancer and rare Mendelian disease [1–4]. Maintaining the security and authenticity of clinical reports containing genetic testing results is an essential component of a clinical laboratory’s workflow as they contain protected health information (PHI) as well as genomic findings that impact healthcare[5,6]. As the continuous rapid advancement of genetic understanding necessitates re-review of previous findings there are often updates to clinical reports, resulting in multiple report ‘versions’ [7,8]. Further, reports can be altered or damaged, even after they have passed beyond the clinical laboratories control. The importance of ensuring that reports that reach patients and clinicians are authentic representations of the most recent and updated versions of those issued from the diagnostic laboratories is therefore an ongoing challenge.

A common solution for this data tracking challenge is for a clinical laboratory to maintain a database of individual report updates [9]. This centralized approach has the advantage of providing complete and immediate data access to information from selected individuals, but requires dedicated staff to maintain centralized compute resources and permission structures for indefinite periods. The central databases are also unable to detect whether reports have been altered subsequent to issue, either maliciously or accidentally. Communication of report authenticity to individuals who are external to the clinical laboratory who do not have database access, including patients, clinicians and auditors is also challenge.[6] Another approach is to issue signed certificates alongside reports; however, this shifts the problem to tracking the certificates themselves and does not provide a mechanism for determining whether a signed report is the most recent.

Here, we present the **A**uthenticated **R**esources in Chained **B**l**o**ck **R**egistry (ARBoR), a simple and efficient approach that addresses the difficulties of monitoring report authenticity by providing a record of cryptographically signed reports, stored in a replicated ledger. This approach augments the secure delivery path between clinical lab and downstream EMR system by providing a durable method to verify the authenticity of files and detect whether newer files are known to exist.

As an alternative to a centralized database, ARBoR employs a file-based ledger that resides in a secure cloud location, written to only by trusted agents and replicated by authorized ARBoR users. By using a block-chaining technique, the ARBoR ledger is verifiable, and the use of strong encryption prohibits the ledger from being modified by malicious actors. Although it has similarities to a “blockchain”, used for example by cryptocurrencies, ARBoR is not a true blockchain as it lacks a decentralized consensus mechanism[10–13] (see discussion). Though these designs are more feature-rich than what we have currently implemented for ARBoR, such blockchain technologies may have broad applicability in the field of genomics[14] and recently, tools like MedRec[15,16] have been implemented to track medical records using blockchain technology. ARBoR instead, balances simplicity and precise tracking to singularly solve the report authenticity problem.[6]

## ARBoR: ARCHITECTURE AND DESIGN

The **A**uthenticated **R**esources in Chained **B**l**o**ck **R**egistry (ARBoR) system (**Figure 1**) implements a replicated ledger of cryptographically-signed clinical reports, enabling the institutions receiving those reports to authenticate and establish the report version. The system consists of three parts: first, the ARBoR Service, which is integrated into the reporting laboratory’s clinical interpretation and reporting pipeline and is responsible for creating a record in a centralized, publicly-readable ledger for the clinical report and related files. Second, the ARBoR Client, which is typically run by an institutional end user as part of the electronic medical record (EMR) ingestion process, receives a replicate of the ledger and uses it to authenticate and validate new clinical reports and related files. Lastly, ARBoRScan (**Figure 2**) is a mobile app for both iOS and Android platforms which also receives a replicate of the ledger that is used by an institutional end-user to verify the authenticity and version of either a printed or Electronic Medical Record (EMR) integrated report before use.

**Figure 1:**
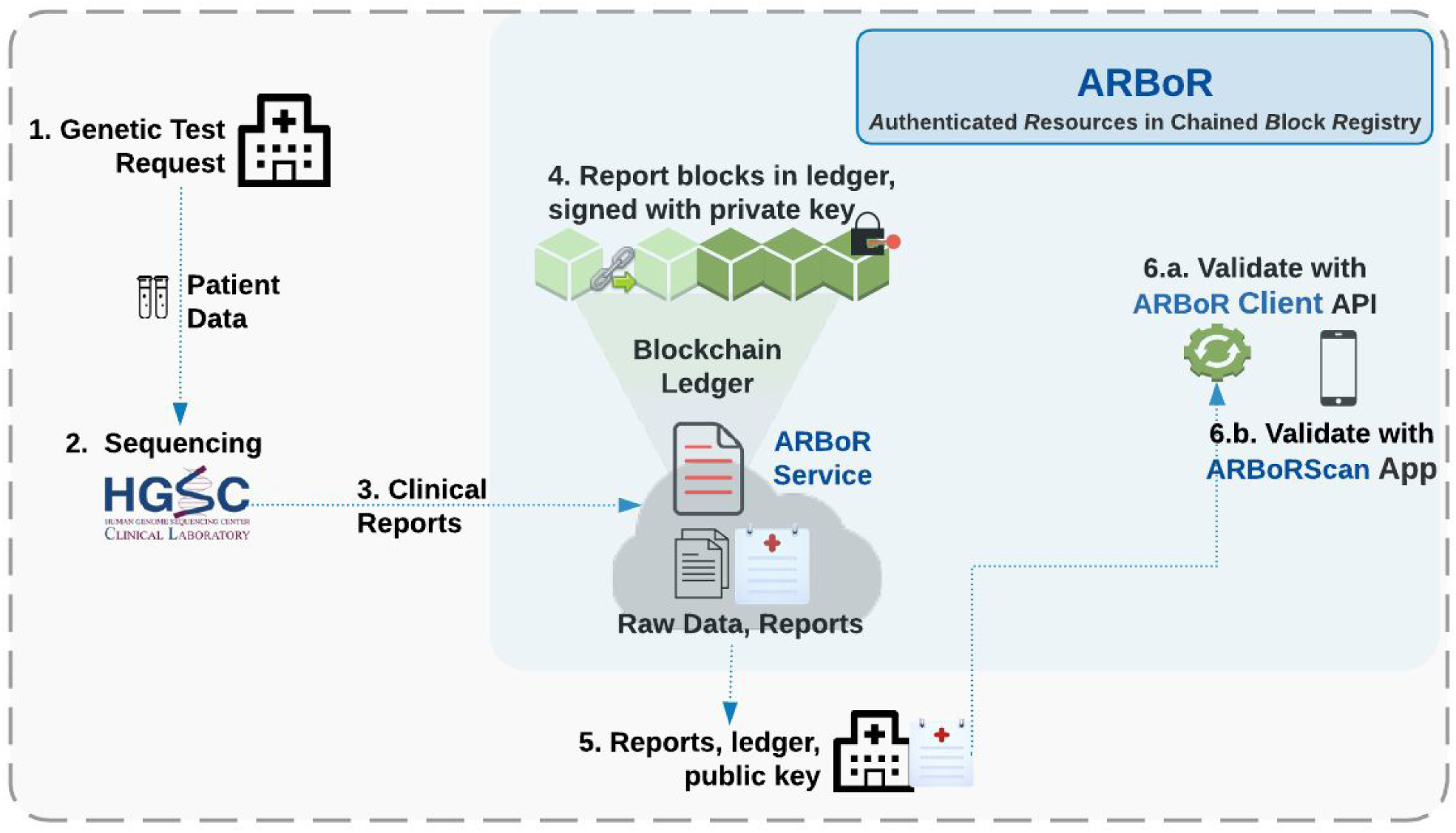
Overall Design of system. The overall ARBoR system consists of three parts: 1: ARBoR Service is integrated into the pipeline of a clinical laboratory. Once the pipeline has generated a clinical report and related files, it uses the ARBoR Service to create and store a record about this file in a public ledger. 2: ARBoR Client is typically run by an institutional end user as part of the ingestion process for new clinical reports and related files. 3: ARBoRScan (Figure 2) is a mobile app for both iOS and Android platforms and is typically run by an end user to fetch metadata about existing reports and check the authenticity and versions of these reports. It also maintains a local copy of the data structure. The primary input is scanning QR codes from existing reports.

**Figure 2:**
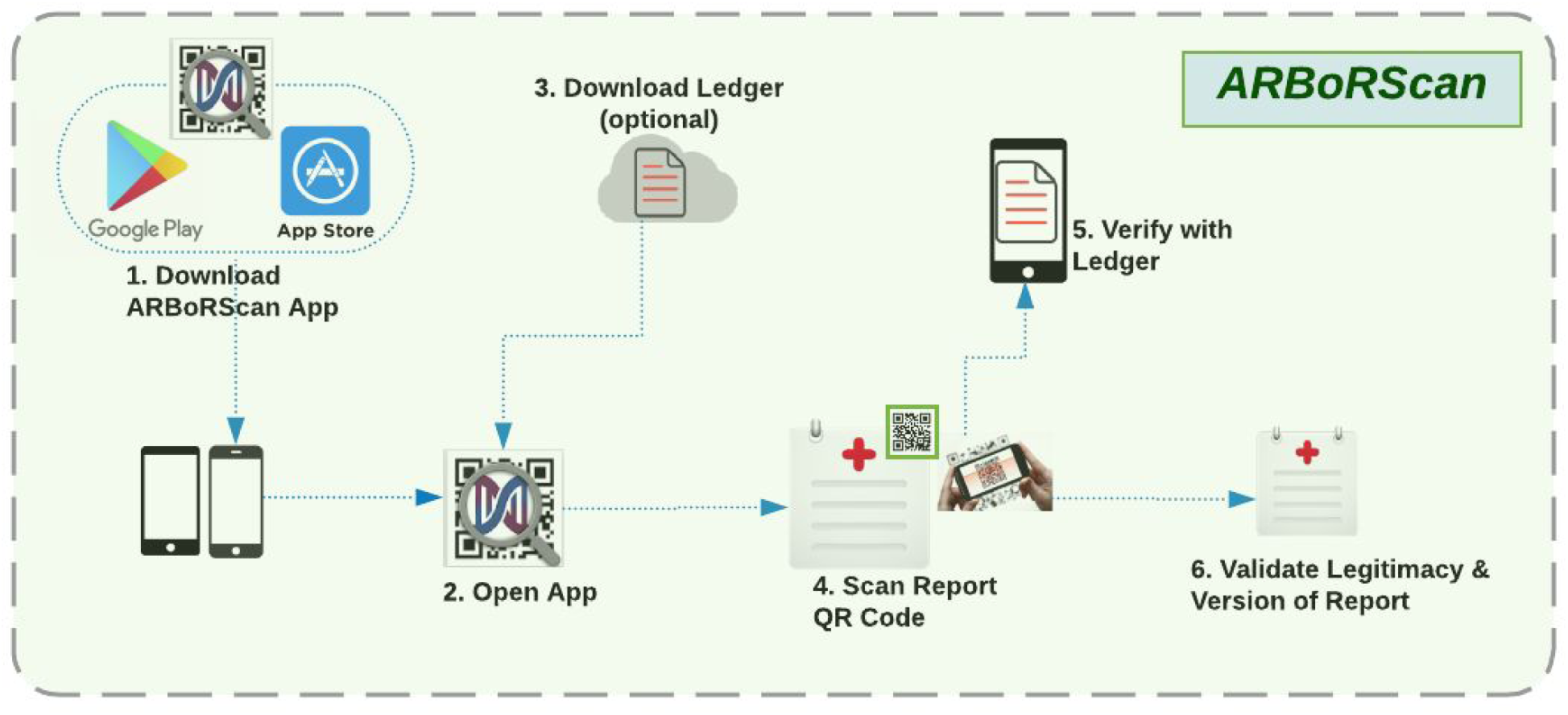
ARBoRScan Process Flow. The mobile app ARBoRScan is available for free download from both iOS and Android App Stores. Accessing the app will automatically check for the latest copy of the ledger; users have the option to download the latest copy of the ledger or use the local version on the device. Users will be able to use this app to scan the QR code (2D barcoded hashed report identifier) on either a printed or EMR integrated report and verify the authenticity and version of the report before use.

The publicly-readable ledger stores records of clinical reports for multiple samples as cryptographically signed blocks. Each block represents a single clinical report and stores metadata about the sample, cryptographically signed contents of the report or file, and block metadata. (**Table 1**). Block metadata consists of a timestamp, the hashed contents of the previous block and the digital signature of the contents of the current block. The previous blockhash is required for every block except the first (the “genesis block”), and subsequent block hashes form a “chain”. Hashing uses the strong SHA3-512[17] algorithm.

**Table 1:**
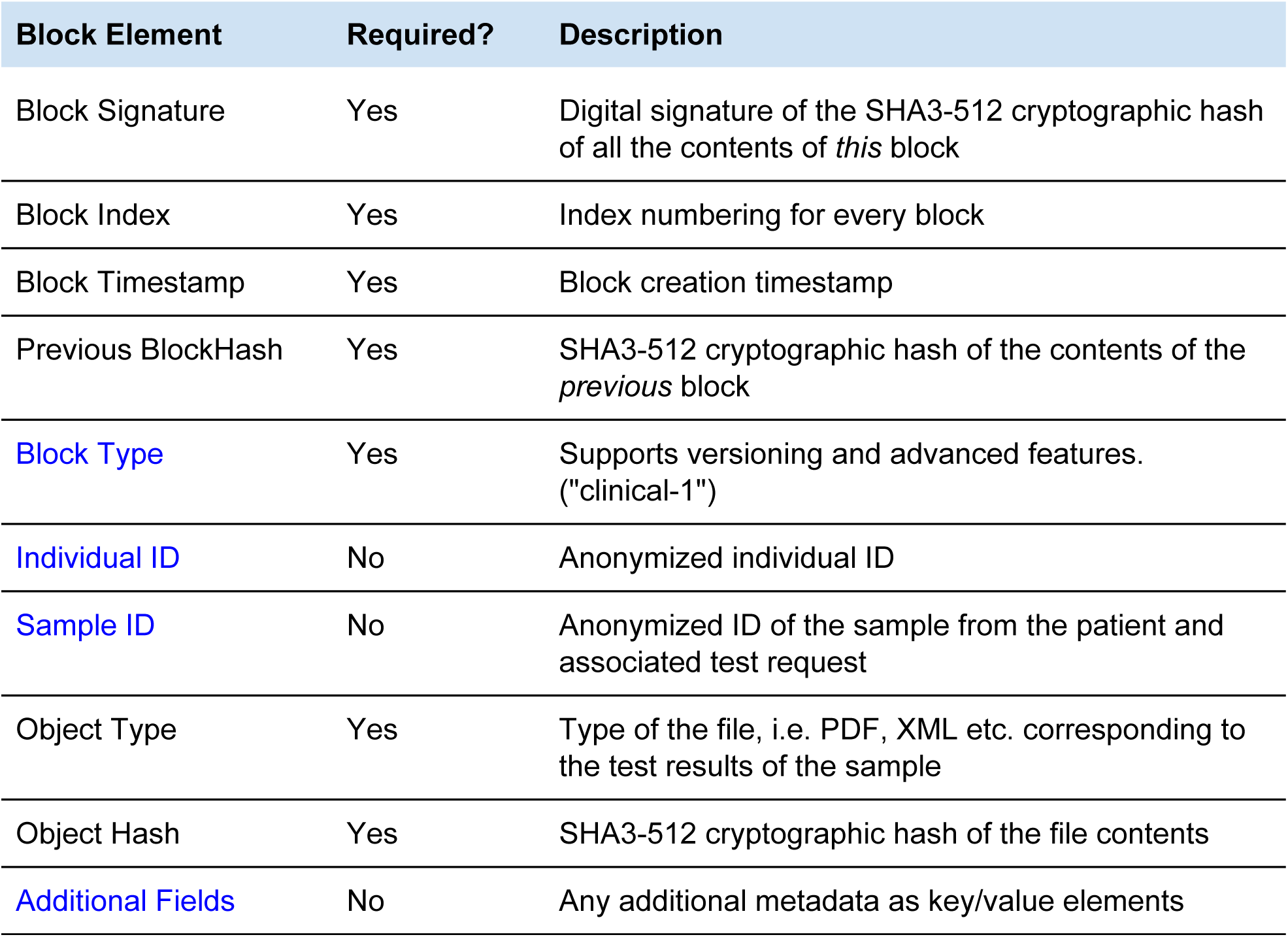
Contents of each block in the ledger. Each block describes a clinical report, and is generated by the ARBoR Service. Input to the Service ranges from the required minimum of a file or folder to file/folders and its associated metadata. Elements in black are auto generated by the Service and elements in blue are parsed and obtained from metadata input. The Object Type allows for the inclusion of multiple types of artefacts and the Block Type allows for the ARBoR software to evolve and add new features without invalidating existing ledgers.

| Block Element | Required? | Description |
| --- | --- | --- |
| Block Signature | Yes | Digital signature of the SHA3-512 cryptographic hash of all the contents of *this* block |
| Block Index | Yes | Index numbering for every block |
| Block Timestamp | Yes | Block creation timestamp |
| Previous BlockHash | Yes | SHA3-512 cryptographic hash of the contents of the *previous* block |
| Block Type | Yes | Supports versioning and advanced features. ("clinical-1") |
| Individual ID | No | Anonymized individual ID |
| Sample ID | No | Anonymized ID of the sample from the patient and associated test request |
| Object Type | Yes | Type of the file, i.e. PDF, XML etc. corresponding to the test results of the sample |
| Object Hash | Yes | SHA3-512 cryptographic hash of the file contents |
| Additional Fields | No | Any additional metadata as key/value elements |

## ARBoR: ENGINEERING AND USE

### Adding New Reports to the Ledger

ARBoR generates one or more public / private key pairs that are specific to a clinical laboratory. The ARBoR Service retains the private keys to sign new blocks. The ARBoR Client and ARBoR App use the associated public keys to authenticate blocks.

To submit a new block to the ledger, the ARBoR Service builds a report block using the private key; the Service verifies the digital signature of the block against the set of trusted public keys, checks the timestamp against the previous entries, and it verifies that the new block contains the block hash of the most recent block in the block chain. This final check could fail due to data corruption, tampering or because the Service needs to fetch an updated copy of the block chain. In that case, the service will traverse the block chain and transmit an error message with the number of missing blocks. If all checks pass, then the new block is appended to the block chain.

### Updating Reports

If a report is updated, a new record is added for that report to the ledger by making a call to the ARBoR Service API. The encrypted ledger entries for prior iterations of the report remain in the ledger, and the new entry is linked to the entry for previous versions of the report. These entries form a ‘chain’ that record the history of the reports for that sample and prevent tampering with the ledger and report history. The ledger can then be distributed along with the report and the public key, usually timed with a data freeze or other bulk release of reports.

### Verifying Digital Copies of Reports

An institutional user with a report file, the ledger, and the laboratory’s public key can use the ARBoR Client software to verify that the report has a valid ledger entry. ARBoR Client compares the signature of the report to the object hash in the ledger to verify the authenticity of a report. The ledger can also be used to check whether a report is the most recent. Periodically, the ARBoRScan app needs to update its local copy of the ledger. To do so, it transmits a RESTful GET request over HTTPS with the hash of the most recent block of which it has knowledge. The ARBoR service will then transmit all blocks that came after the block mentioned in the request. Alternatively, the app can verify the electronic version of a report file by using the Python API tools available on [github link].

### Verifying Printed Reports

To verify a printed report, a user first scans the report’s QR code using ARBoRScan. This app connects to the latest version of the report ledger, extracts the hashed report ID from the QR code, and retrieves the corresponding ledger entry. Once a report has been verified, the user can optionally be directed to a website that contains additional information about that report, or alternatively allows printing of a new version to replace one that has been defaced or damaged. This approach can remains effective even subsequent to the conclusion of the project without requiring elaborate resources for maintenance and upkeep.

### Security

Taken together, the set of stringent checks described above create a secure system. First, security of ARBoR Service’s addition of new blocks to the ledger is guaranteed by requiring that a new block be created only by a clinical laboratory holding a valid private key and requiring that transactions occur over https. Next, neither the clinical pipeline nor the ARBoR Client fully trust the ARBoR Service. By checking the current ledger in the Service against previously known blocks and known public keys both the pipeline and the client are in a position to detect alterations in the external behavior of the Service. Furthermore, the ARBoR Client checks all incoming blocks against its set of public keys and validation rules. While the clinical reports themselves are delivered outside of ARBoR, as depicted in the above sections, ARBoR ensures the authenticity and integrity of the reports and files delivered, thereby allowing the downstream EMR to reject untrusted files.

## DISCUSSION

We introduced the ARBoR system for tracking and versioning clinical reports. It produces an encrypted ledger which can be replicated and provided to clinical partners for use in verifying their copies of clinical reports. ARBoR consists of three components: ARBoR Service is employed to produce the ledger, whereas ARBoR Scan and ARBoR Client own replicated copies of the ledger and use it to verify reports. This system allows us to create a tamper evident, easily verifiable, secure record of clinical report histories.

The ledger is centralized in that only trusted instances of the ARBoR Service may write to it. Future iterations of ARBoR may function as a decentralized system, if for example we need to support multiple clinical laboratories issuing reports. This decentralization would require communication between ARBoR Service instances, which would need a mechanism to agree upon the latest ledger. These include functionality and complexity not required for our use case.

These were not in the current implementation but can be incorporated into future versions. A natural preliminary extension would be a ‘private blockchain’ design[18] which allows only trusted agents to write to the ledger and simplifies the consensus mechanism. Further, ARBoR Client and ARBoR Scan are designed to function without an active instance of ARBoR Service. At the end of a project, the ARBoR Service instance for that project can be retired, with the final ledger being a deliverable of the project. So long as the ARBoR Client and ARBoR Scan instances have received the final ledger, no centralized infrastructure need be maintained.

Although initially developed for tracking clinical reports, this approach is extensible to any file type. The Human Genome Sequencing Center’s Clinical lab at Baylor College of Medicine frequently produces other exported data deliverables (e.g. vcf, bam and cram files) and there are use cases where it is desired to securely link additional metadata (e.g. quality control metrics) to these files in perpetuity. The ability to track these files with ARBoR is a future area of development. Another extension that we are exploring is to use ARBoR to track the actual report delivery transactions. This would require that report delivery be recorded by the ARBoR Service as an entry in the ledger as an element in the chain with the report that was delivered.

An important benefit to using a chained block structure for the ledger is that the ledger as a whole can be re-validated at any time. Validating the ledger consists of starting with the most recent block and then following the chain, checking that hashing each block gives the expected value recorded in the following block. Additionally, since each block is digitally signed, it is possible to detect if fraudulent block(s) have been added. Finally, the timestamps of the blocks should be in chronological order.

We have deployed this technology for our eMERGE project, where clinical reports are issued to clinical sites around the country. Reports are distributed in XML and pdf formats; using ARBoR, both report formats can be authenticated and checked whether they contain the latest information. this approach provides a durable method for tracking all of our deliverables, ensuring their authenticity and data integrity with a complete audit trail. To date, this approach has aided the data management of 15,205 reports.

## CONCLUSIONS

ARBoR provides clinical laboratories with a simple and efficient means for tracking clinical reports and provides a distributable ledger that can travel with reports to their recipients. This ledger provides a means of validating clinical reports, that persists well beyond the lifespan of a project. We have successfully applied this system in the context of the eMERGE clinical sequencing project. ARBoR code is available on GitHub at https://github.com/BCM-HGSC/ARBoR and the mobile app ARBoRScan is available for iOS at https://goo.gl/QZXpqg and for Android at https://goo.gl/cLdKB8.

